# Distinct neural moderators of resilience and vulnerability confer heterogeneous outcomes following early-life adversity

**DOI:** 10.1101/2025.09.16.676492

**Authors:** Huaxin Fan, Benjamin Becker, Jie Zhang

## Abstract

Early-life adversity (ELA) increases risk for psychiatric disorders, but outcomes are highly variable. Using multimodal neuroimaging and computational moderation in large adolescent cohorts, we systematically identify brain features that buffer or amplify effects of three ELA subtypes—familial interpersonal, non-familial interpersonal, and non-interpersonal adversity—on transdiagnostic psychopathology. Multimodal neural moderators were subtype– and pathology dimension-specific, revealing distinct neurobiological mechanisms moderating heterogeneous outcomes. Protective features clustered in limbic, sensory integration, and regulatory circuits (amygdala, parietal cortex, anterior cingulate), while vulnerability features concentrated in frontotemporal circuits. Individual features and a new Relative Resilience Index (RRI)—reflecting the individual balance between protective and vulnerability features across the whole-brain—prospectively predicted psychopathological progression over two years, linking brain signatures to future mental health trajectories. Findings establish the brain as dynamic moderator of distinct adversity effects and introduce a system-level marker for risk stratification, advancing mechanistic precision in youth mental health and guiding early intervention.

## Introduction

Early-life adversity (ELA) is a major determinant of psychiatric disorders and suicide risk,^1^ and exerts substantial societal and economic costs.^2^ However, individuals exposed to similar types of adversity during early life frequently display strikingly divergent life-course and clinical outcomes, suggesting that biological or psychological factors can buffer or exacerbate effects of ELA.^3–7^ In line with contemporary resilience research—which shifts the focus from mechanisms of disease vulnerability to those supporting adaptive stress resistance—these individual differences highlight the need to identify both resilience and vulnerability factors.^8^ Such moderators are critical for understanding heterogenous risk trajectories and for developing personalized prevention strategies.^9–11^ Adolescence—a developmental window characterized by brain maturation and heightened vulnerability to psychopathology—offers an exceptional window when distinct developmental trajectories emerge, determining whether individuals exposed to ELA will experience enduring adverse consequences or exhibit adaptive resistance.^12,13^

Moderation models provide a powerful framework to determine factors that influence the strength or direction of ELA’s impact on mental health.^4,14,15^ Personality traits have been extensively examined as moderators and personality profiles that attenuate adverse effects are considered protective, while those that amplify them confer vulnerability.^3,4,14,15^ While a range of environmental and psychological moderators have been examined in relation to ELA and psychopathological outcomes,^16^ a growing body of research is now focusing on the brain as a key source of individual variability. Advances in neuroimaging have facilitated investigations into the underlying mechanisms, and extensive evidence has linked ELA to alterations in brain regions involved in emotion regulation, executive control, reward processing, social cognition, and threat response.^17–20^ Crucially, emerging evidence is shifting the paradigm from considering the brain merely as passive endpoint of adversity, to recognizing it as active moderator that can shape the impact of ELA on psychopathology. Certain neural characteristics may function as moderators—buffering or amplifying psychological outcomes—rather than serving as passive markers of early stress. For instance, increased functional connectivity between the amygdala and insula, elevated nodal efficiency, and stronger whole-cortex structure–function coupling have been shown to amplify adversity-related risk, indicating potential neural vulnerability factors.^5,21,22^ In contrast, greater activation to positive socio-emotional stimuli in the left pallidum, increased engagement of the anterior prefrontal cortex during emotion regulation, and a connectome-based brain marker involving frontoparietal and subcortical regions have been linked to attenuate such risk, suggesting potential neural protective factors.^23–25^ Shifting the focus from ELA-induced brain changes to brain-based moderators of ELA’s effects could allow prediction of an elevated risk of detrimental long-term effects of ELA and reveal novel targets for early intervention to enhance resilience and mitigate long-term risk. However, most existing studies have relied on hypothesis-driven approaches, and systematic, data-driven identification of neural moderators remains limited. Moreover, the predominance of single-modality imaging has yielded fragmented insights, constraining our understanding of how integrated neural systems confer vulnerability or protection to ELA. Multimodal neuroimaging offers a promising path forward by integrating complementary measures of brain structure and function, enabling a more comprehensive characterization of individual differences in susceptibility and resilience to ELA.^26^

ELA is a heterogeneous construct. Traditional frameworks, such as the adverse childhood experiences (ACEs) model, emphasize interpersonal adversity within the family.^27^ However, ecological systems theory highlights that developmental outcomes emerge from multiple interacting environmental layers.^28^ In adolescence, adversity increasingly extends beyond the family to include peers and broader community contexts, which in turn impact on the brain and psychopathological trajectories.^12,29,30^ Therefore, a refined taxonomy capturing both interpersonal and broader environmental contexts is essential. Emerging evidence demonstrates that interpersonal and non-interpersonal adversities constitute distinct dimensions, each engaging dissociable psychological and neurobiological pathways.^31–37^ Interpersonal adversity—defined by intentional harm or threat from others—undermines fundamental aspects of trust and compassion, leading to enduring disruptions in social adjustment.^20,31,32^ This adversity has been linked to accelerated development in fronto-limbic cicruits and more severe psychopathological outcomes than non-interpersonal adversity.^31,33,36,37^ In contrast, non-interpersonal adversity (e.g., accidents, natural disasters, and bereavement)-which is characterized by the absence of intentionality – has been associated with delayed development in temporal-limbic regions.^32,33,36^ Further distinctions within interpersonal adversity are also of key significance. Familial interpersonal adversity (e.g., violence occurring within the home) may exert fundamental distinct developmental consequences to non-familial interpersonal adversity (e.g., violence occurring outside the home).^38–40^ In line with prior work,^31^ we adopt a tripartite classification of adversity—familial interpersonal, non-familial interpersonal, and non-interpersonal—to provide a more nuanced framework for investigating whether distinct brain moderators exhibit subtype-specific effects.

The consequences of early adversity frequently transcend traditional diagnostic categories. Rather than conferring the risk of specific categorical disorders, ELA’s impact is more accurately reflected in transdiagnostic dimensions—such as internalizing, externalizing, and the general psychopathology (p) factor—that span multiple diagnostic categories based on underlying behavioral and neural alterations.^41,42^ Given that these dimensions show distinct neurobiological specificity,^43–45^ individual differences in psychological outcomes may reflect variations in brain features that modulate environmental sensitivity across these symptom domains. However, it remains unclear whether and how specific neural characteristics confer susceptibility or resilience by differentially moderating the effects of distinct adversity types across multiple psychopathological dimensions. Addressing this gap is essential to elucidate the neural pathways shaping mental health trajectories after early adversity, with direct implications for targeted prevention and intervention.

Here, we leverage a large neurodevelopmental dataset to systematically investigate whether specific brain features moderate associations between diverse forms of ELA and transdiagnostic psychopathology in adolescence. We focus on three adversity subtypes—familial interpersonal, non-familial interpersonal, and non-interpersonal adversity—and three broad symptom dimensions: p factor, internalizing, and externalizing. By systematically mapping brain–adversity interactions across symptom domains and adversity subtypes, our goal is to identify candidate neural markers that drive individual differences in vulnerability or resilience to distinct environmental challenges, providing a critical step towards personalized prediction and intervention in youth mental health.

## Methods

### Participants

Data were obtained from the Adolescent Brain Cognitive Development (ABCD) Study (Release 3.0), comprising approximately 11,878 participants enrolled at ages 9–10 years. Written informed consent and assent were obtained from parents and children, respectively. The study protocol was approved by the Institutional Review Board at the University of California, San Diego, which oversees ethical compliance.^46^ Full details on study procedures are available from the ABCD Study consortium.^47–49^

### Early-life adversity

We examined three broad categories of ELA: familial interpersonal adversity (FIA), non-familial interpersonal adversity (NFIA), and non-interpersonal adversity (NIA). According to prior literature,^32,34,35,37^ interpersonal adversity (IA) includes physical, emotional, and sexual abuse; physical and emotional neglect; community violence; bullying; and war, while NIA encompasses exposures such as natural disasters, accidents, and sudden major losses. To operationalize these constructs within the ABCD dataset, we identified relevant items from validated instruments used in previous ABCD studies focusing on baseline ELA.^47,50–53^ Specifically, FIA was defined as adversity originating from family members, including physical, emotional, and sexual abuse; physical and emotional neglect; and domestic violence. NFIA included adversity from members outside family, such as physical, emotional, and sexual abuse; bullying; school safety concerns; other community violence; and war. NIA comprised natural disaster, accidents, and sudden losses. For each subtype, item-level responses were aggregated to compute a separate total score, yielding one adversity score per subtype for each participant. Detailed descriptions of the specific measures and scoring procedures are provided in the Supplementary Table S1.

### Psychopathology

Consistent with prior work,^47^ a general psychopathology (P) factor and orthogonal internalizing (INT) and externalizing (EXT) factors were derived from baseline parent-reported Child Behavior Checklist (CBCL) data. A bifactor model was fitted to eight CBCL subscales—Withdrawn, Somatic Complaints, Anxious/Depressed, Social Problems, Thought Problems, Attention Problems, Delinquent Behavior, and Aggressive Behavior. The general P factor captured shared variance across all subscales; the EXT-specific factor loaded on Delinquent and Aggressive Behavior; and the INT-specific factor loaded on Withdrawn, Somatic Complaints, and Anxious/Depressed. Factor scores were extracted and used in subsequent analyses. Model fit was acceptable by conventional criteria (χ^2^ = 1362.436, *df* = 16, *p* < 0.001; RMSEA = 0.084; CFI = 0.973; TLI = 0.954; SRMR = 0.028), supporting both statistical adequacy and theoretical interpretability.

In addition, psychiatric diagnoses at baseline and at the 2-year follow-up were obtained via parent-report using the computerized Kiddie Schedule for Affective Disorders and Schizophrenia (KSADS) for DSM-5. For the present analysis, lifetime diagnoses were coded dichotomously (0 = no diagnosis, 1 = any diagnosis) at each time point, regardless of specificity. Change in diagnostic status between baseline and follow-up was used to index clinical progression.

### Multimodal neuroimaging acquisition and processing

Detailed descriptions of the ABCD Study neuroimaging acquisition protocols and scanner parameters are available elsewhere,^48^ as are image processing and analysis methods.^49^ For the present analyses, we used curated and tabulated neuroimaging data provided by the ABCD consortium. Participants were excluded from analyses if their scans failed visual inspection or did not meet FreeSurfer quality control standards (e.g., imgincl_t1w_include = 1).

We included multimodal brain measures derived from structural MRI, diffusion imaging, and resting-state and task-based functional MRI. Structural MRI (SMRI) included 354 measures capturing cortical thickness, surface area, volume, sulcal depth, and T1-weighted contrast across 68 cortical regions defined by the Desikan atlas, as well as the volume of 14 subcortical regions (bilateral thalamus, caudate, putamen, pallidum, hippocampus, amygdala, and nucleus accumbens). Diffusion tensor imaging (DTI) comprised 234 measures, including fractional anisotropy (FA) and mean diffusivity (MD) derived from a full-shell model across 35 major white matter tracts,^54^ pericortical white matter regions, and subcortical structures. Resting-state fMRI (RSFMRI) included 273 measures of functional connectivity (FC) within and between 13 cortical networks (based on the Gordon parcellation), as well as between each network and subcortical regions. Task-based fMRI measures were derived from three tasks. The Monetary Incentive Delay (MID) task provided 328 activation measures across cortical and subcortical regions for four contrasts: large reward vs. neutral anticipation, large loss vs. neutral anticipation, positive vs. negative reward feedback, and positive vs. negative loss feedback. The Stop Signal Task (SST) contributed 164 activation measures based on correct stop vs. correct go and incorrect stop vs. correct go contrasts. Finally, the emotional n-back task (ENBACK) yielded 164 activation measures for negative vs. neutral and positive vs. neutral face conditions.

### Relative resilience index (RRI)

To quantify an individual’s overall resilience, we computed a Relative Resilience Index (RRI) integrating adversity exposure with the presence of resilient and vulnerable brain features, as follows:

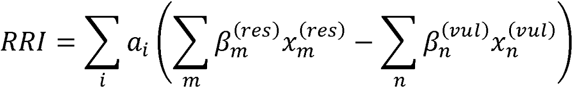

Here, *a_i_* indicates the presence (1) or absence (0) of adversity *i*,. Each resilient feature 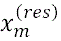 and vulnerable feature 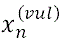 was trichotomized based on their distributions following prior research that used ±1 standard deviation (SD) cutoffs to define high– and low-trait resilience groups.^55^ Specifically, values exceeding one SD above the mean were coded as 1 (presence), those below one SD as −1 (absence), and values within ±1 SD as 0 (neutral). The weights 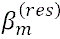 and 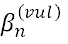 represent standardized coefficients from the interaction terms involving the respective features in moderation analyses, reflecting their relative contribution.

## Statistical analyses

### Moderation analysis

Moderation models provide a robust framework to examine factors that influence the strength or direction of ELA’s effects on mental health.^4,14,15^ The model can be expressed as:

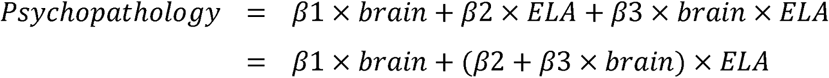

In this formulation, the marginal effect of *ELA* on *Psychopathology* is given by *β*2 + *β*3 × *brain*. Traits that attenuate adverse effects are considered protective, while those that amplify them confer vulnerability.^3,4,14,15^ Accordingly, a positive *β*3 indicates that larger values of the brain feature increase the negative impact of *ELA* on *Psychopathology*, suggesting a vulnerability factor; conversely, a negative *β*3 suggests the brain feature reduces this impact, indicating a protective factor.

This study employed linear mixed-effects models to assess how brain features moderate the associations between three types of ELA (FIA, NFIA, and NIA) and psychopathology dimensions (P, EXT, or INT). Each model incorporated one brain feature, the three ELA types as main effects, their brain × ELA interaction terms, and relevant covariates—including age, sex, race, family income, parental education, BMI, MRI manufacturer, and modality-specific neuroimaging covariates (intracranial volume for T1-weighted data and mean framewise displacement for other modalities). Random intercepts were specified for family ID nested within acquisition site ID to account for hierarchical data structure, consistent with previous study.^43^ To control for multiple comparisons, Bonferroni correction was applied separately for each combination of adversity type, neuroimaging modality, and psychopathology dimension. Interaction terms with Bonferroni-adjusted *p* < 0.05 were considered significant; standardized regression coefficients (*β*) with 95% confidence intervals (CI) were reported alongside their corresponding brain features.

To provide a more intuitive understanding of the moderating effects of the identified brain features, we performed simple slope analysis and marginal effects analysis. For simple slope analysis, we estimated the association between ELA and psychopathology at representative levels of the brain feature (mean, meanLJ+LJ1 SD, and meanLJ−LJ1 SD). For marginal effects analysis, we evaluated the conditional effect of ELA on psychopathology across the continuous range of brain feature values, generating confidence intervals to visualize the regions where the effect was significant.

To assess model robustness, the dataset was divided into training (80%) and test (20%) sets, with ten-fold cross-validation performed on the training data. Model performance was evaluated by calculating Pearson’s correlation coefficients between predicted and observed psychopathology scores in the test set.

### Ordinal Logistic Regression

To further examine whether the identified protective and vulnerability-related neural factors, as well as individual’s overall resilience (i.e., the RRI), are associated with clinical progression, we employed ordinal logistic regression. Clinical progression was treated as an ordinal variable with three categories: improved (scored −1), unchanged (scored 0), and worsened (scored 1), making ordinal logistic regression an appropriate method for modeling these ordered outcomes.^56^ To ensure robustness, we applied ten-fold cross-validation. When examining the links between each brain feature and disease progression, analyses were restricted to individuals exposed to the specific adversity corresponding to that feature. For these individuals, each brain feature was binarized based on prior research that used ±1 SD cutoffs to define high– and low-trait resilience groups (coded as 1 if > mean + 1 SD, and 0 if < mean − 1 SD).^55^ We hypothesized that RRI and resilient features would be negatively associated with clinical deterioration, while vulnerability features would show positive associations. To test these directional hypotheses, one-sided *p*-values were derived from the *z*-statistics of the ordinal logistic regression models. Odds ratios (OR), 95% CI, and one-sided *p*-values are reported.

## Results

### Identification and distribution of protective and vulnerability brain features

We defined neural markers that attenuate the association between ELA and psychopathology as protective, and those that strengthen it as vulnerability factors. Screening 1,517 candidate brain features using moderation models, we identified 57 that remained significant after Bonferroni correction (Fig. 1a), including 34 with negative *brain* × *ELA* interaction coefficients (standardized *β* range: –0.086 to –0.048; consistent with a protective role) and 23 with positive coefficients (standardized β range: 0.047 to 0.073; consistent with a vulnerability role). These features spanned multiple modalities (Fig. 1b), with their proportion varying across modalities (calculated relative to the total number of candidate features within each modality): functional activity during reward processing (MID-task) showed the highest proportion (5.8%), white matter connectivity (DTI) the lowest (1.3%), and SMRI, RSFMRI, SST, and ENBACK each approximately 4%.

**Fig. 1:**
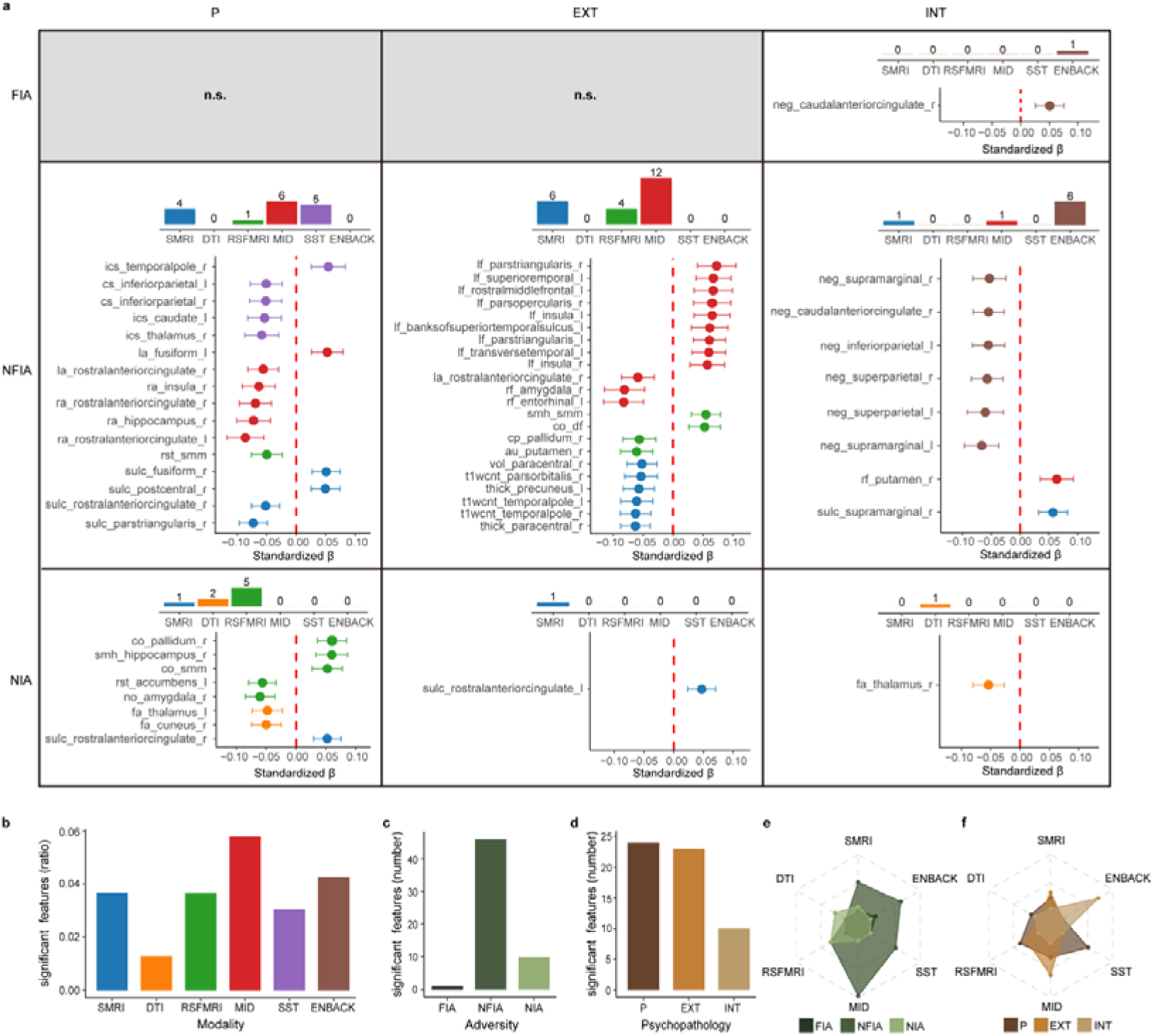
Multimodal neural moderators of the ELA–psychopathology relationship. **a**, Point-and-line plot showing standardized *brain* × *ELA* interaction coefficients and 95% CI for significant features across ELA–psychopathology combinations. Negative coefficients indicate protective features, and positive coefficients indicate vulnerability features. Bar plots show the number of significant features in each modality for all ELA–psychopathology combinations. **b–d**, Distribution of significant features across modalities (**b**), ELA types (**c**), and psychopathology dimensions (**d**). **e–f**, Distribution of significant neural features across modalities for each ELA type (**e**) and psychopathology dimension (**f**). sulc, sulcus depth. vol, cortical volume. t1wcnt, T1-weighted contrast. thick, cortical thickness. fa, fractional anisotropy. rst, retrosplenial-temporal network. smm, sensorimotor mouth network. co, cingulo-opercular network. smh, sensorimotor hand network. no, none network (including regions from orbitofrontal and temporal cortex). df, default network. au, auditory network. ics, incorrect stop. cs, correct stop. la, large loss versus neutral anticipation. ra, large reward versus neutral anticipation. lf, positive versus negative loss feedback. rf, positive versus negative reward feedback. neg, negative versus neutral face. r, right hemisphere. l, left hemisphere. n.s., not significant.

In line with our hypotheses the number and specific distribution of neural features also differed across ELA types and psychopathology dimensions (Fig. 1c-f). Across ELA types, 46 features moderated NFIA, 10 moderated NIA, and only one moderated FIA. NIA-related features were restricted to structural (DTI) and resting-state (RSFMRI) modalities, whereas NFIA-related features spanned structural, resting-state, and all three task-based modalities. For psychopathology dimensions, more than 20 features moderated the effect of ELA on P and EXT, whereas 10 moderated its effect on INT. For P, all modalities except ENBACK contained a moderate and balanced number of features, with SST features observed exclusively for P. EXT-related features were limited to SMRI, RSFMRI, and MID, with MID contributing the largest number (12 features). INT involved SMRI, DTI, and MID, each with a single feature, whereas ENBACK contributed seven features, all observed only for INT.

Within each modality, contrasts revealed variation in both the number of significant features and the direction of *brain* × *ELA* interaction coefficients across Within each modality, contrasts revealed variation in both the number of significant ELA–psychopathology combinations (Fig. 1a). In structural modalities, SMRI contrasts were concentrated in sulcal depth, cortical thickness, and T1-weighted contrast, with sulcal depth showing both negative and positive coefficients, and thickness and T1-weighted contrast observed only with negative coefficients in NFIA–EXT associations. For DTI, only a few FA measures in pericortical white matter regions and subcortical structures were significant, showing negative coefficients, whereas major tracts and MD showed no significant interactions. RSFMRI features were concentrated in motor-related circuits, including the sensorimotor cortex and basal ganglia. For task-based modalities, SST contrasts of incorrect stop vs correct go and correct stop vs correct go involved a similar number of significant features in NFIA–P associations, predominantly with negative coefficients. MID features were primarily observed in P-related anticipation contrasts with negative coefficients and EXT-related feedback contrasts with positive coefficients, all within NFIA associations. ENBACK features were specific to negative-face contrasts and mostly observed in NFIA–INT associations with negative coefficients.

Different types of ELA were associated with distinct neural moderators of psychopathology, highlighting the heterogeneity of their neurobiological pathways (Fig. 2). For FIA, no significant negative *brain* × *ELA* interactions were observed; the only significant positive effect was activation of the right caudal anterior cingulate cortex (ACC) during negative versus neutral face processing (standardized *β* = 0.051, 95% CI [0.026, 0.076], *p* = 6.9×10^-5^; Fig. 2a, right). By contrast, NFIA exhibited widespread neural moderation, with both negative and positive coefficients spanning multiple domains. Negative coefficients involved bilateral parietal cortex, right inferior frontal gyrus, limbic regions (bilateral ACC, temporal pole, right insula, hippocampus, amygdala, left entorhinal cortex), and motor-related circuits (sensorimotor network, right thalamus, bilateral basal ganglia) together with their FC with the cingulo-parietal, retrosplenial-temporal, and auditory networks. Clusters of effects were evident, such as four features in the right rostral ACC and two each in the left inferior parietal lobule and right paracentral gyrus. The strongest negative effect was located in the activation of left rostral ACC during large reward versus neutral anticipation (standardized β = –0.086, 95% CI [-0.118, –0.055], *p* = 7.7×10^-8^; Fig. 2b, left). Positive coefficients were concentrated in bilateral prefrontal and temporal cortices, insula, motor circuits (sensorimotor network, right putamen), and FC between the default and cingulo-opercular networks, with the strongest positive effect in the activation of right pars triangularis during positive versus negative loss feedback (standardized β = 0.073, 95% CI [0.040, 0.105], *p* = 1.1×10^-5^; Fig. 2b, right). For NIA, negative coefficients were concentrated in subcortical and limbic regions, including bilateral thalamus, left nucleus accumbens, right amygdala, the none network including regions from orbitofrontal and temporal cortex, the retrosplenial-temporal network, and right precuneus, with the strongest effect in FC between the none network and right amygdala (standardized β = –0.060, 95% CI [-0.085, –0.036], *p* = 1.8×10^-6^; Fig. 2c, left). Positive coefficients involved bilateral rostral ACC, motor-related circuits (sensorimotor network, right pallidum) and their FC with the cingulo-opercular network and right hippocampus, with the strongest effect in the FC between cingulo-opercular network and right pallidum (standardized β = 0.060, 95% CI [0.036, 0.084], *p* = 1.3×10^-6^; Fig. 2c, right).

**Fig. 2:**
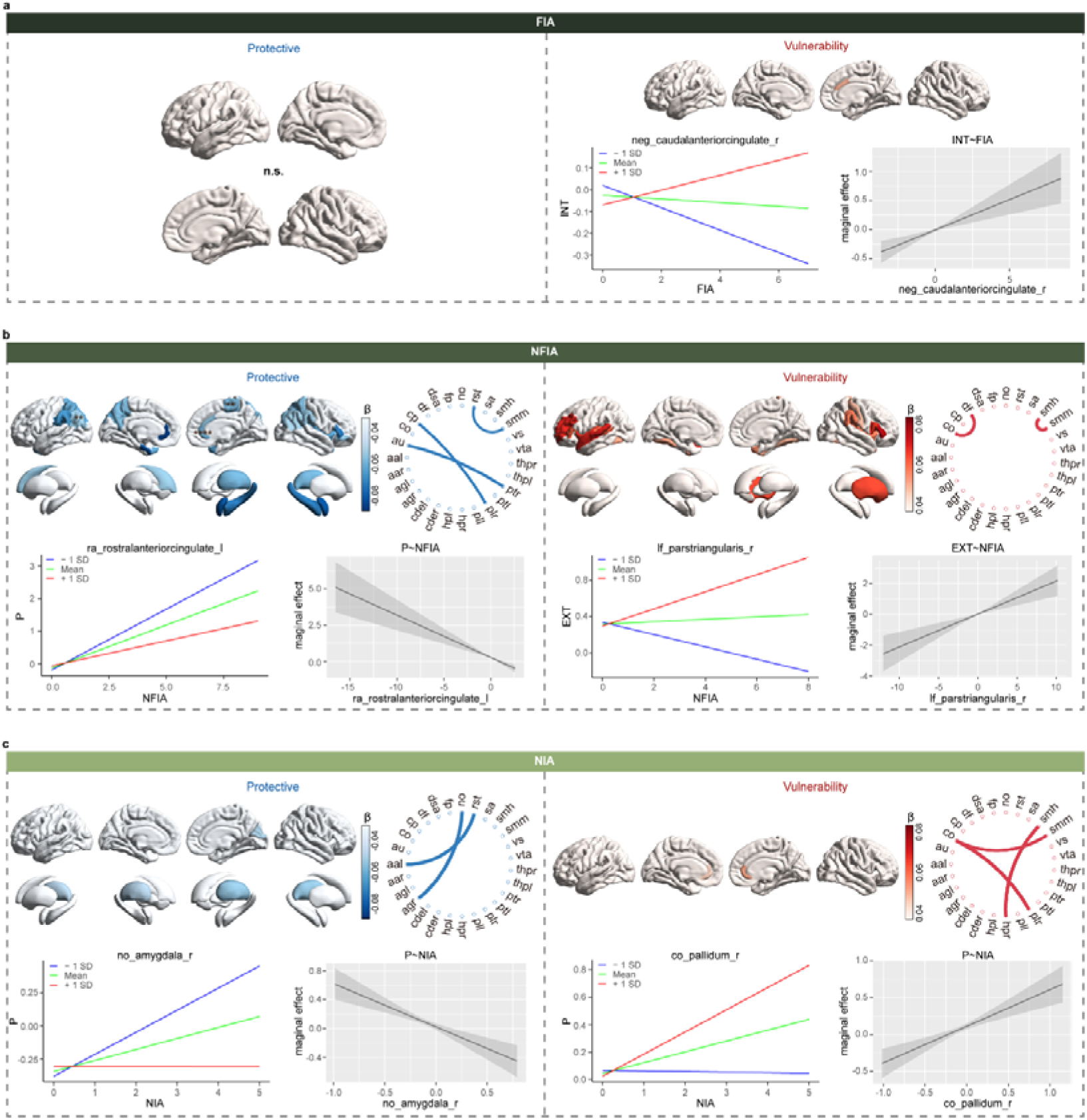
Spatial distribution of protective and vulnerability brain features and their key moderation effects on ELA types. **a-c**, Distinct spatial distribution neural moderators were observed across FIA (**a**), NFIA (**b**), and NIA (**c**). β represents the mean standardized coefficient of *brain × ELA* interaction across all significant features within each brain region. Negative coefficients indicated protective features, and positive coefficients indicated vulnerability features. The number of asterisks on each brain region corresponds to the number of significant features; regions with color but no asterisk represent a single feature. Selected features with the strongest negative and positive coefficients are highlighted with simple slope and marginal effects analyses for each ELA type. Simple slope plots show the ELA–psychopathology association at low, medium, and high levels of the brain feature. Marginal effects plots show how the conditional effect of ELA on psychopathology varies across the continuous range of the brain feature, with shaded areas representing 95% CI. au, auditory network. co, cingulo-opercular network. cp, cingulo-parietal network. df, default network. dsa, dorsal attention network. fp, frontal-parietal network. no, none network. rst, retrosplenial-temporal network. sa, salience network. smh, sensorimotor hand network. smm, sensorimotor mouth network. vs, visual network. vta, ventral attention network. thp, thalamus proper. pt, putamen. pl, pallidum. hp, hippocampus. cde, caudate. ag, amygdala. aa, nucleus accumbens. r, right hemisphere. l, left hemisphere.

Distinct spatial patterns also characterized neural moderators of different psychopathology dimensions (Fig. 3). For P, negative coefficients spanned multiple domains, including bilateral inferior parietal lobule, right inferior frontal gyrus, limbic regions (bilateral ACC, right insula, hippocampus, amygdala, left nucleus accumbens, and the none network), and motor-related circuits (sensorimotor network, bilateral thalamus, left caudate), with three significant features in the right rostral ACC and the strongest effect in the activation of left rostral ACC during large reward versus neutral anticipation (standardized β = –0.086, 95% CI [-0.118, –0.055], *p* = 7.7×10^-8^; Fig. 3a, left). Positive coefficients were located in bilateral temporal cortex, right ACC, motor circuits (sensorimotor network, right pallidum) and their FC with the cingulo-parietal network and right hippocampus, with the strongest effect in FC between cingulo-opercular network and right pallidum (standardized β = 0.060, 95% CI [0.036, 0.084], *p* = 1.3×10LJLJ; Fig. 3a, right). For EXT, negative coefficients involved left precuneus, right inferior frontal gyrus, limbic regions (left entorhinal cortex, right ACC, right amygdala), and motor-related circuits (right paracentral lobule, basal ganglia) including their FC with cingulo-parietal and auditory networks; two features were significant in the right paracentral lobule, with the strongest effect in the activation of left entorhinal cortex during positive versus negative reward feedback (standardized β = –0.083, 95% CI [-0.116, –0.049], *p* = 1.1×10^-6^; Fig. 3b, left). Positive coefficients were more spatially clustered, involving bilateral prefrontal cortex, insula, left superior temporal gyrus, and FC within the sensorimotor network and between the default and cingulo-opercular networks, with the strongest effect in activation of right pars triangularis during positive versus negative loss feedback (standardized β = 0.073, 95% CI [0.040, 0.105], *p* = 1.1×10^-5^; Fig. 3b, right). For INT, negative coefficients were concentrated in bilateral parietal cortex, right thalamus, and right ACC, with the strongest effect in the activation of left supramarginal cortex during negative versus neutral face processing (standardized β = –0.067, 95% CI [-0.097, –0.037], *p* = 1.2×10^-5^; Fig. 3c, left). Positive coefficients were located in right supramarginal gyrus, ACC, and putamen, with the strongest effect in right putamen activation during positive versus negative reward feedback (standardized β = 0.064, 95% CI [0.035, 0.092], *p* = 1.5×10^-5^; Fig. 3c, right). Supplementary Fig. S1–S3 show the simple slope and marginal effects plots for each brain feature that significantly moderated the effect of each ELA type on P, EXT, and INT.

**Fig. 3:**
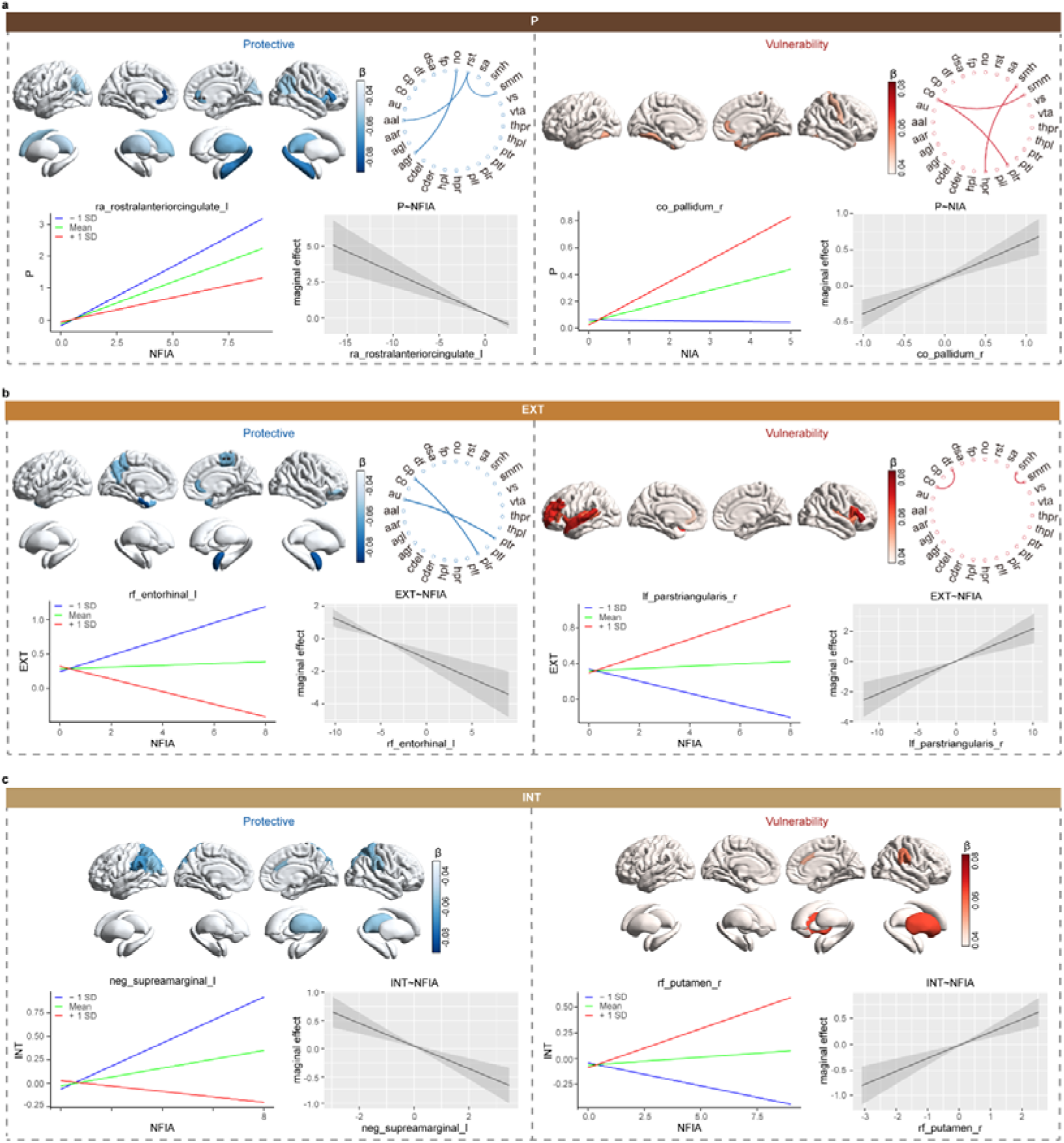
Spatial distribution of protective and vulnerability brain features and their key moderation effects on psychopathology dimensions. **a-c**, Distinct spatial distribution neural moderators were observed across P (**a**), EXT (**b**), and INT (**c**). β represents the mean standardized coefficient of *brain × ELA* interaction across all significant features within each brain region. Negative coefficients indicated protective features, and positive coefficients indicated vulnerability features. The number of asterisks on each brain region corresponds to the number of significant features; regions with color but no asterisk represent a single feature. Selected features with the strongest negative and positive coefficients are highlighted with simple slope and marginal effects analyses for each psychopathology dimension. Simple slope plots show the ELA–psychopathology association at low, medium, and high levels of the brain feature. Marginal effects plots show how the conditional effect of ELA on psychopathology varies across the continuous range of the brain feature, with shaded areas representing 95% CI.

Together, these findings show that different forms of ELA and psychopathology dimensions are moderated by separable neural patterns. Some associations were widely distributed across multiple domains, while others were spatially clustered within specific circuits.

Most brain features exerted significant moderating effects in only a single ELA–psychopathology combination. Notably, three features in the right ACC showed significant modulation across multiple combinations, with consistent or opposing effects (Fig. 4). Activation of the right rostral ACC during large loss versus neutral anticipation negatively moderated NFIA’s effects on both P and EXT. The sulcus depth of the same region negatively moderated NFIA’s effect on P but positively moderated NIA’s effect on P. Likewise, activation of the right caudal ACC during negative versus neutral face negatively moderated NFIA’s effect on INT while positively moderating FIA’s effect on INT.

**Fig. 4:**
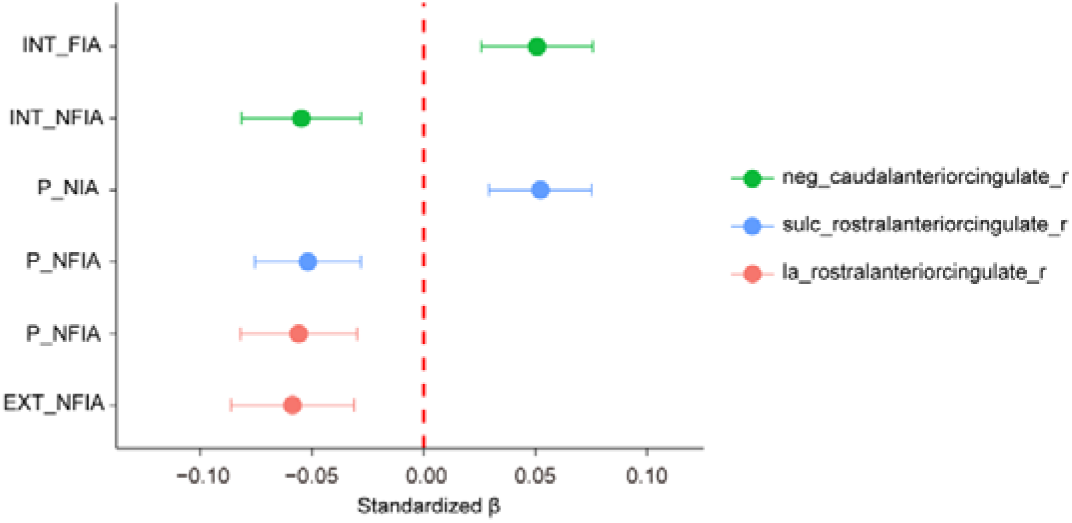
Brain features showing consistent or opposing moderation effects across multiple ELA–psychopathology combinations.

### Predictive performance of brain features and integrated RRI for longitudinal outcomes

To assess the reliability of the moderation models, we applied the significant models from the training set to the test set and computed correlations between predicted and observed psychopathology scores. Across all modality–ELA–psychopathology combinations, correlation coefficients ranged from 0.37 to 0.87 (Fig. 5), demonstrating moderate to strong reliability.

**Fig. 5:**
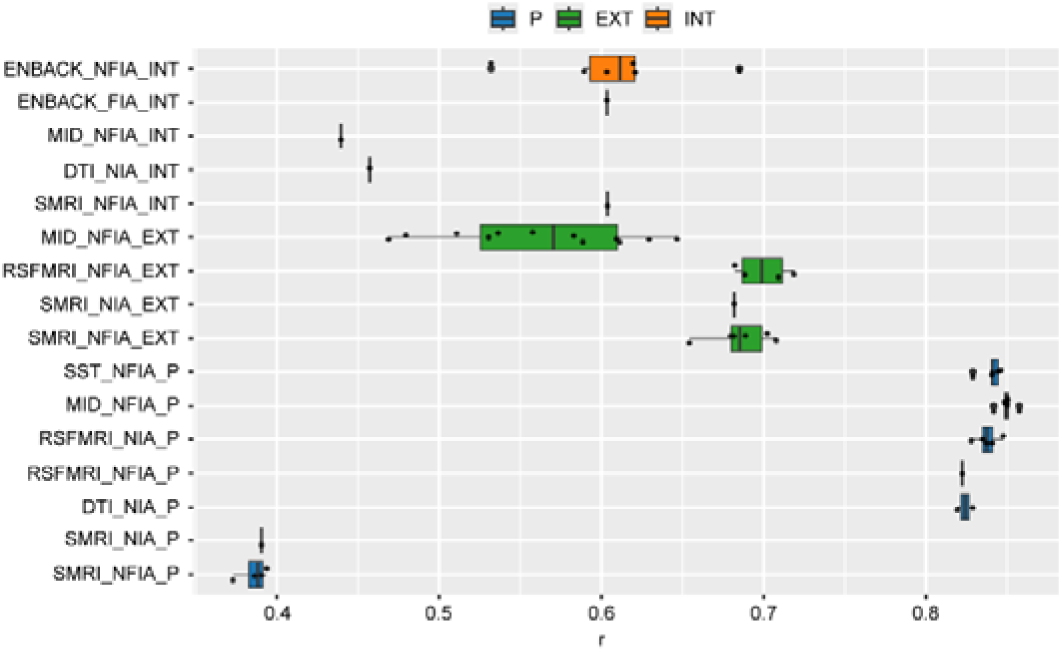
Test-set correlations between predicted and observed psychopathology values for brain features. Each point represents a single feature within a modality–ELA–psychopathology combination, and boxplots summarize the distribution of correlations across features (range: 0.37–0.87), illustrating the predictive performance of the moderation models.

To further evaluate whether the putative protective and vulnerability brain features identified in the moderation analyses could prospectively predict clinical progression, we conducted ordinal logistic regression analyses. Specifically, we tested whether individuals exposed to a given type of ELA, who also exhibited the corresponding protective or vulnerability neural feature, showed a reduced or elevated risk of overall psychiatric deterioration two years later. Among the 57 candidate features, only six significantly predicted longitudinal outcomes in the hypothesized direction. Three features—which in the moderation analyses showed negative interaction coefficients—activity of the right rostral ACC during anticipation of large loss versus neutral (OR 0.27, 95% CI [0.09, 0.81], *p* = 0.010, one-sided), cortical thickness of the right paracentral gyrus (OR 0.39, 95% CI [0.13, 1.07], *p* = 0.033, one-sided), and activity of the right inferior parietal cortex during correct stop versus correct go (OR 0.76, 95% CI [0.55, 1.04], *p* = 0.045, one-sided)—were associated with reduced likelihood of clinical deterioration. In contrast, three features—which in the moderation analyses showed positive interaction coefficients—activity of the left transverse temporal cortex (OR 6.18, 95% CI [1.08, 35.45], *p* = 0.021, one-sided) and left insula (OR 1.42, 95% CI [0.99, 2.04], *p* = 0.028, one-sided) during positive versus negative loss feedback, and FC between sensorimotor hand network and right hippocampus (OR 1.30, 95% CI [0.98, 1.71], *p* = 0.034, one-sided)—were linked to increased risk of deterioration.

Given that individuals may experience multiple types of ELA and exhibit multiple protective or vulnerability brain features across the whole brain (Fig. 6), we derived a composite measure of relative resilience—the RRI—to quantify the overall balance of protective versus vulnerability features at the whole-brain level. Ordinal logistic regression analyses showed that RRI predicted reduced risk of overall psychiatric deterioration two years later (OR 0.69, 95% CI [0.49, 0.96], *p* = 0.013, one-sided).

**Fig. 6:**
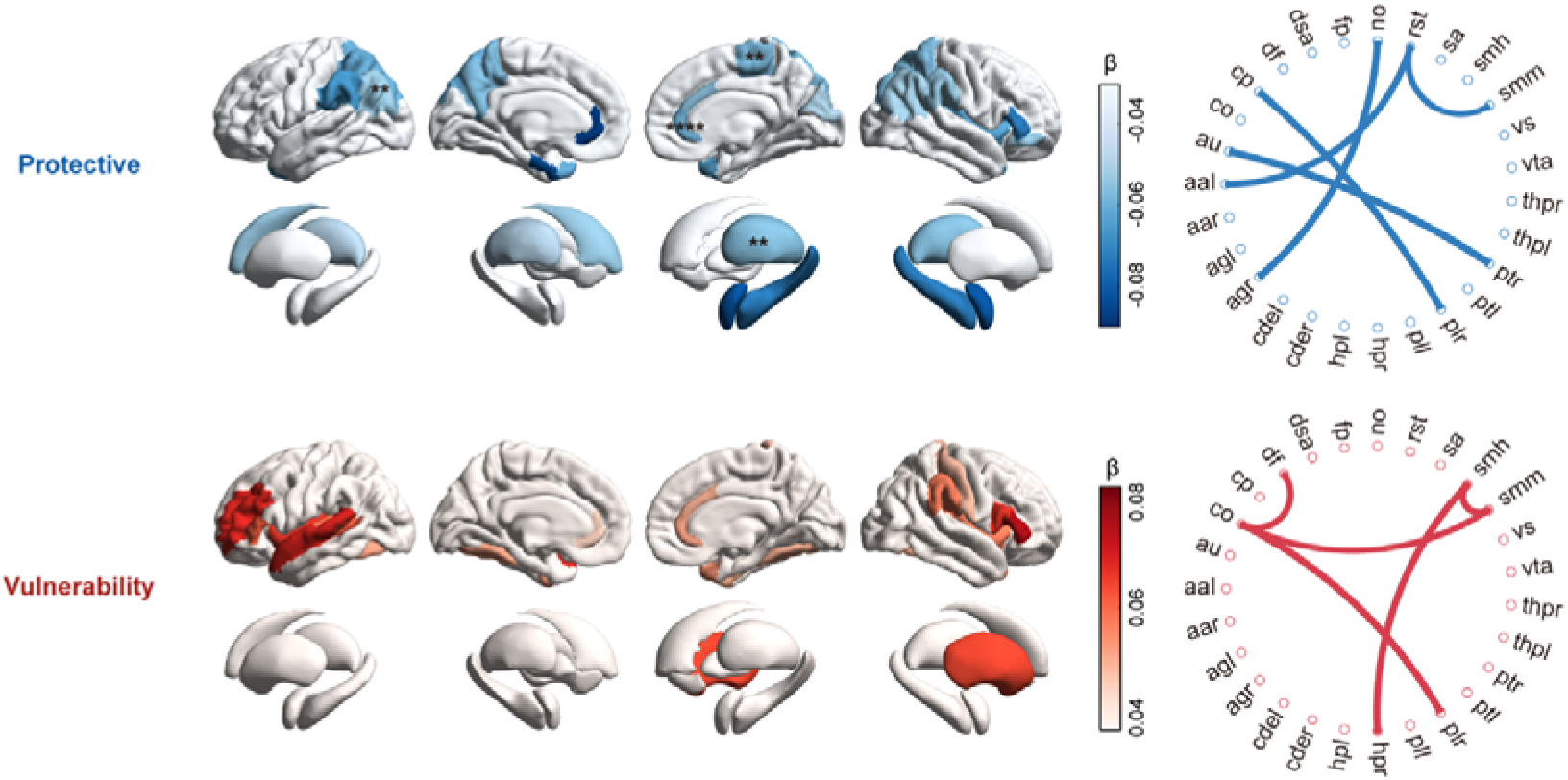
Spatial distribution of integrated protective and vulnerability brain features across ELA types and psychopathology dimensions. β represents the mean standardized coefficient of *brain × ELA* interaction across all significant features within each brain region. Negative coefficients indicated protective features, and positive coefficients indicated vulnerability features. The number of asterisks on each brain region corresponds to the number of significant features; regions with color but no asterisk represent a single feature.

## Discussion

In this large adolescent cohort, leveraging a comprehensive multimodal, data-driven framework, we systematically identified 57 robust neural moderators of ELA from 1,517 candidate brain features, including 34 conferring protection and 23 conferring vulnerability on the effect of ELA on later psychopathology. Combining the moderation approach with ELA subtypes and a dimensional psychopathology framework allowed us to determine distinct neural moderators that are specific to both ELA subtype and psychopathology dimensions. Notably, protective brain features were concentrated in limbic, sensory integration, and regulatory circuits (including amygdala, insula, hippocampus, parietal cortex, ACC), while vulnerability features were predominately localized in frontal and temporal systems, particularly during threat and loss avoidance contexts. These findings underscore the utility of moderation models in identifying brain features that specifically buffer or amplify the impact of adversity on psychopathology. Importantly some neural features, and particularly our RRI—aggregating protective and vulnerability features across the whole brain to quantify each individual’s balance between resilience and vulnerability across different adversity contexts—predicted reduced risk of psychiatric deterioration over the following two years. These findings present predictive evidence that the brain serves not only as passive residue of adversity, but as an active regulator that can shape the trajectories of ELA. These findings support a conceptual shift from viewing the brain solely as an “outcome” of adversity to recognizing it as a dynamic moderator influencing vulnerability and resilience in an ELA context and system-specific manner. Strikingly, FIA was only moderated by a single neural marker, while NIA and NFIA where moderated by considerably more factors (10 or 46, respectively) putatively reflecting the fundamental detrimental impact of adversity within the family. Unlike prior hypothesis-driven, single-modality studies,^5,21–23,25^ our cross-modality and cross-task analyses indicate that protection and vulnerability are not confined to a single circuit but reflect coordinated, multi-system organization. The distribution of significant effects across modalities points to potential pathways: reward-processing features (MID) accounted for the largest proportion (5.8%), white-matter structure (DTI) the smallest (1.3%), with SMRI, RSFMRI, SST, and ENBACK each contributing about 4%. This pattern supports evidence that the reward system can moderate ELA’s effects on psychopathology,^57,58^ and underscores the value of multimodal integration—such as the RRI—for understanding the link between ELA and psychopathology.

ELA is a well-established risk factor for a wide range of psychiatric symptoms and suicide risk,^1^ and it imposes substantial economic burdens on society.^2^ Understanding the regulatory mechanisms that buffer or amplify the impact of ELA thus carries significant clinical relevance for prevention and early intervention.^8–10^ While genetic studies have identified some moderators,^59^ genetic features are not directly modifiable. Our findings highlight the brain’s unique role as a dynamic moderator: protective features attenuate the ELA–psychopathology association, whereas vulnerability features amplify it, forming a bidirectional regulatory mechanism. Given the brain’s plasticity, these moderators represent actionable targets for cognitive behavioral therapy, noninvasive brain stimulation, or pharmacological interventions. By integrating protective and vulnerability features across the whole brain, the RRI provides a systems-level measure of resilience that can identify individuals at elevated risk. Collectively, these results advance mechanistic understanding of ELA-related psychopathology and offer translational potential for clinical applications.

The neural circuits underlying moderation differed by ELA type. NFIA showed the most widespread effects, likely reflecting the challenge posed by intentional social threats to higher-order cognitive–emotional integration networks.^36^ Protective features were concentrated in limbic system (ACC, entorhinal cortex, insula, amygdala, hippocampus), thalamus, frontoparietal control regions, and basal ganglia–sensorimotor loops. Together, these interconnected systems may support resilience and adaptive stress regulation through their critical role in a range of social, affective and reward related functions including cognitive control and self-regulation, autonomic and emotion regulation, social approach and interpersonal functions, and reward and motivation.^60–66^ They also underscore the beneficial role of an adaptive allostatic interoception system in moderating the effects of NFIA. This system enables the brain to anticipate and regulate bodily responses via neural, neuroendocrine, and immune pathways, dynamically adjusting heart rate, hormone secretion, and metabolism, thereby reducing allostatic load and promoting efficient stress management.^10^ In contrast, vulnerability features were most pronounced in the frontotemporal cortices, particularly during successful lossLJavoidance feedback, suggesting heightened sensitivity to self-relevant and socially threatening cues.^67,68^ Increased coupling between the default mode and cingulo-opercular networks—typically anti-correlated—also characterized vulnerability, reflecting a failure to appropriately segregate internally– and externally-directed processing and an inclination toward excessive rumination.^69,70^ For NIA moderation, relatively few frontal, temporal, or parietal regions were engaged. Instead, effects were concentrated within the allostatic interoception system, suggesting that non-intentional events influence mental health primarily by disrupting bodily regulation and interoceptive processing, rather than through higher-order social-cognitive pathways. The greater involvement of brain regions in NFIA compared with NIA is consistent with prior evidence.^71^ By contrast, FIA exhibited limited modulation, with only hyperactivation of the right caudal ACC in response to negative faces emerging as a vulnerability marker. While this concurs with the ACC’s role in cognitive and emotion regulation it also suggests that the neurodevelopmental impact of adverse experiences within the family may be so profound that they override the adaptive capacity of the brain, effectively diminishing the potential for neural systems to buffer or counteract the impact of familial adversity^72^ Overall, different ELA types act via distinct neural circuits, providing a clear framework for understanding their unique effects on mental health.

Moderation effects also varied by psychopathology dimension. P-factor moderators spanned control, reward, and sensorimotor regions, consistent with the view that general psychopathology engages broad brain networks,^73^ while SST features specifically underscored the moderating role of conflict monitoring and inhibitory control.^74^ EXT moderators were also widespread, spanning frontal and parietal regions, motor-related areas, and reward circuits, reflecting the influence of impulsivity and reward sensitivity.^75^ Although both P and EXT were strongly moderated by MID reward-related features, these effects showed dimension specificity, reward/loss specificity, and stage-dependent divergence. During reward anticipation, protective features were prominent, marked by increased activation in the ACC, insula, and hippocampus, primarily affecting the P factor. This suggests that proactive control and strategic preparation may buffer ELA’s adverse impact on general psychopathology.^76,77^ By contrast, during successful loss-avoidance feedback, vulnerability features were more pronounced, with heightened activation in the prefrontal cortex, insula, and superior temporal gyrus, primarily affecting EXT. This likely reflects excessive neural reactivity or insufficient regulation of rumination,^78^ increasing risk for behavioral impulsivity and externalizing symptoms.^79^ INT moderators were more localized, concentrated in the parietal cortex, ACC, thalamus, and striatum, underscoring the central role of emotion–attention integration circuits in anxiety– and depression-related internalizing problems.^80^ Notably, ENBACK features were selectively linked to INT during negative face processing, underscoring the specificity of affective processing—particularly the attentional bias toward negative stimuli—in internalizing psychopathology.^81^

Among all moderators, the ACC stood out as particularly salient. Its effects recurred across ELA types and psychopathology dimensions, with functional and structural features showing opposing influences. For example, greater sulcal depth of the right rostral ACC conferred protection for P under NFIA but vulnerability under NIA, while right caudal ACC activation to negative faces bidirectionally moderated INT. These findings suggest that the ACC, as a hub for valuation, reappraisal, and conflict monitoring,^72^ serves as a key node within multi-system regulatory networks, with modulatory effects shaped by adversity context, task stage, and structural constraints.

Our findings align with the RDoC framework,^82^ emphasizing transdiagnostic, multi-domain, and environment-integrated approaches, while highlighting a mechanism-focused understanding of psychopathology. The multimodal moderators we identified map closely onto RDoC’s core functional domains—cognitive control, reward, negative valence, social processes, and sensorimotor systems—and exert effects across ELA–psychopathology combinations. Whereas numerous prior studies have documented cross-domain brain network changes following adversity and their impact on mental health,^17–19,83,84^ our results suggest that specific network configurations may actively determine an individual’s resilience or vulnerability to particular forms of adversity. This mechanism-based perspective reframes the brain from a passive outcome of adversity to an active regulator of its effects, complementing prior work and providing a novel framework for understanding individual differences.

Individuals often experience multiple forms of ELA and exhibit diverse protective and vulnerability brain features. Given that these features may accumulate to collectively confer resilience,^10^ we developed the RRI to integrate their effects into a whole-brain index. By aggregating across ELA–psychopathology combinations, the RRI quantifies each individual’s balance between resilience and vulnerability at the neural systems level. This framework aligns with the broader approach of integrating protective and risk factors to characterize brain states,^85^ and resonates with the One Health concept, which unites external exposome and internal biological factors to predict wellbeing.^86,87^ Compared with single-region metrics, the RRI captures convergence across tasks and modalities, reducing the influence of task-specific or incidental effects and providing a more stable neural phenotype. Although most single features showed limited predictive power at follow-up—likely due to complex nonlinear relationships between symptoms and diagnoses^88^—the integrated RRI significantly predicted reduced risk of overall clinical deterioration over two years (OR = 0.69), demonstrating that it captures synergistic multi-system signals and holds promise as a prospective predictive tool.

Adolescence represents a critical window of heightened neural plasticity, during which cortical gyrification and myelination increase—key hallmarks of brain maturation.^89,90^ Importantly, the developmental trajectory of the prefrontal cortex often lags behind earlier-maturing subcortical regions such as the amygdala and hippocampus,^13,91^ creating variability in prefrontal–subcortical maturation across individuals. According to this mismatch framework, individuals whose prefrontal regions develop relatively faster may be better able to regulate subcortical activity, thereby exhibiting neural resilience to adversity-related disruption. Consistent with this notion, our results identified increased sulcus depth and white matter contrast in the right ACC and inferior frontal gyrus as protective neural features associated with NFIA, reflecting aspects of prefrontal maturation that may enhance regulation of downstream circuits. These regional moderators provide a basis for the RRI, which can help identify adolescents at high risk and guide personalized interventions tailored to their adversity exposures and psychopathology profiles—such as enhancing cognitive control, reward sensitivity, or emotion regulation—interventions in these domains during this stage can yield long-lasting benefits.^10,11^ During intervention, neuroimaging could monitor protective versus vulnerability brain activity in real time,^11^ potentially augmented by noninvasive brain stimulation to suppress vulnerability circuits and strengthen protective ones.^92,93^ This approach, aligned with the RDoC framework, integrates neuropsychological phenotypes to create a closed-loop pathway from phenotype identification to individualized intervention. As a whole-brain integrative measure, the RRI captures an individual’s overall risk or resilience, complementing specific single-brain features that can serve as actionable targets for intervention, thereby providing a bridge for translating statistical findings into clinically actionable targets. Applied during adolescence, this dual approach may identify those at high risk, enhance neural resilience, reduce vulnerability, and generate positive cascading effects in coping with adversity and psychiatric risk,^94^ establishing a neurobiological foundation for early and targeted interventions.

Methodologically, the study’s strengths include its data-driven approach, large sample size, multimodal imaging, cross-task evaluation, stringent Bonferroni correction, and robust train–test reliability (r = 0.37–0.87). Longitudinal follow-up further enhances external validity and translational potential. Limitations should be noted. First, although we identified protective and vulnerability features and constructed the RRI, the temporal ordering between brain features and adversity exposures remains unclear Establishing causal inference will require more stringent longitudinal or interventional study designs.^95^ Second, while we adjusted for covariates such as age and sex, we did not examine sex-specific differences in neural moderators, and findings are limited to children aged 9–10 years; future longitudinal studies should track developmental trajectories across sexes and ages to inform tailored interventions.^10^ Third, although we integrated multimodal data, we did not investigate how covariance across modalities may jointly moderate adversity–psychopathology associations—an important direction for future work.^96^

In conclusion, by identifying neural moderators of resilience and vulnerability across ELA types and psychopathology dimensions using moderation models and integrating them into the RRI, this study reveals how the brain actively shapes individual sensitivity to adversity. This framework predicts adolescent mental health outcomes and provides a neurobiological basis for precision prevention and intervention, highlighting the value of moderation models in uncovering protective and vulnerability brain features and offering a new mechanistic account of individual differences in mental health trajectories.

## Acknowledgements

We are grateful to Dr. Zhaowen Liu and Dr. Xinran Wu for their valuable suggestions, and to Dr. Yu Liu for technical support. J.Z. was supported by STI2030-Major Projects 2021ZD0200204 and Key Laboratory of Computational Neuroscience and Brain-Inspired Intelligence (Fudan University), Ministry of Education, China. B.B. was supported by National Natural Science Foundation of China (82271583), National Key Research and Development Program of China (2018YFA0701400), and start-up and collaborative seed grants from the University of Hong Kong. H.F. was supported by China Scholarship Council (202406100214).

## Author Contributions

H.F. conceived and designed the study, conducted data analyses, wrote the first draft of the manuscript. H.F., B.B., and J.Z. contributed to critical revision of the report for important intellectual content. All authors have read and approved the final version of the manuscript. B.B., and J.Z. were responsible for the final decision to submit for publication.

## Data Availability

The data that support the findings of this study can be accessed from ABCD 3.0 Data Release at https://nda.nih.gov/study.html?id=901.

