## Supplementary materials for "Distinct neural moderators of resilience and vulnerability confer heterogeneous outcomes following early-life adversity"

### Supplementary Tables

| Variable | Description and Value Notes | Coding | Name of Instrument & Short Name |
| --- | --- | --- | --- |
| **Familial Interpersonal Adversity (Range: 0-8)** | | | |
| ***Emotional Abuse***  *[A ‘1’ on this item contributed a point to the score of Interpersonal Early Adversity (Family)]* | | | |
| ksads_ptsd_raw_765_p | A family member threatened to kill your child.  0 = No; 1 = Yes | - | ABCD Parent Diagnostic Interview for DSM-5 (KSADS) Traumatic Events.  abcd_ptsd01 |
| ***Physical Abuse***  *[A ‘1’ on either item contributed a point to the score of Interpersonal Early Adversity (Family)]* | | | |
| ksads_ptsd_raw_762_p | Shot, stabbed, or beaten brutally by a grown up in the home.  0 = No; 1 = Yes | - | ABCD Parent Diagnostic Interview for DSM-5 (KSADS) Traumatic Events.  abcd_ptsd01 |
| ksads_ptsd_raw_763_p | Beaten to the point of having bruises by a grown up in the home.  0 = No; 1 = Yes | - | ABCD Parent Diagnostic Interview for DSM-5 (KSADS) Traumatic Events.  abcd_ptsd01 |
| ***Sexual Abuse***  *[A ‘1’ on this item contributed a point to the score of Interpersonal Early Adversity (Family)]* | | | |
| ksads_ptsd_raw_767_p | A grown up in the home touched your child in their privates, had your child touch their privates, or did other sexual things to your child.  0 = No; 1 = Yes | - | ABCD Parent Diagnostic Interview for DSM-5 (KSADS) Traumatic Events.  abcd_ptsd01 |
| ***Domestic Violence***  *[To contribute a point to the score of Interpersonal Early Adversity (Family), the response to the item ‘ksads_ptsd_raw_766_p’ should be coded as ‘1’, or at least one response to other items should be coded as ‘1’]* | | | |
| ksads_ptsd_raw_766_p | Witness the grownups in the home push, shove or hit one another.  0 = No; 1 = Yes | - | ABCD Parent Diagnostic Interview for DSM-5 (KSADS) Traumatic Events.  abcd_ptsd01 |
| fes_youth_q1 | We fight a lot in our family.  1 = True; 0 = False | - | ABCD Youth Family Environment Scale-Family Conflict Subscale Modified from PhenX (FES)  abcd_fes01 |
| fes_youth_q2 | Family members rarely become openly angry.  0 = True; 1 = False | - | ABCD Youth Family Environment Scale-Family Conflict Subscale Modified from PhenX (FES)  abcd_fes01 |
| fes_youth_q3 | Family members sometimes get so angry they throw things.  1 = True; 0 = False | - | ABCD Youth Family Environment Scale-Family Conflict Subscale Modified from PhenX (FES)  abcd_fes01 |
| fes_youth_q4 | Family members hardly ever lose their tempers.  0 = True; 1 = False | - | ABCD Youth Family Environment Scale-Family Conflict Subscale Modified from PhenX (FES)  abcd_fes01 |
| fes_youth_q5 | Family members often criticize each other.  1 = True; 0 = False | - | ABCD Youth Family Environment Scale-Family Conflict Subscale Modified from PhenX (FES)  abcd_fes01 |
| fes_youth_q6 | Family members sometimes hit each other.  1 = True; 0 = False | - | ABCD Youth Family Environment Scale-Family Conflict Subscale Modified from PhenX (FES)  abcd_fes01 |
| fes_youth_q7 | If there's a disagreement in our family, we try hard to smooth things over and keep the peace.  0 = True; 1 = False | - | ABCD Youth Family Environment Scale-Family Conflict Subscale Modified from PhenX (FES)  abcd_fes01 |
| fes_youth_q8 | Family members often try to one-up or outdo each other.  1 = True; 0 = False | - | ABCD Youth Family Environment Scale-Family Conflict Subscale Modified from PhenX (FES)  abcd_fes01 |
| fes_youth_q9 | In our family, we believe you don't ever get anywhere by raising your voice.  0 = True; 1 = False | - | ABCD Youth Family Environment Scale-Family Conflict Subscale Modified from PhenX (FES)  abcd_fes01 |
| ***Emotional neglect***  *[To contribute a point to the score of Interpersonal Early Adversity (Family), at least one response should be coded as ‘1’]* | | | |
| crpbi_parent1_y | First caregiver (caregiver participating in study/completing protocol). Makes me feel better after talking over my worries with him/her.  1 = Not like him/her; 2 = Somewhat like him/her; 3 = A lot like him/her | 1 = Not like him/her;  0 = otherwise | ABCD Children's Report of Parental Behavioral Inventory.  crpbi01 |
| crpbi_parent2_y | First caregiver (caregiver participating in study/completing protocol). Smiles at me very often.  1 = Not like him/her; 2 = Somewhat like him/her; 3 = A lot like him/her | 1 = Not like him/her;  0 = otherwise | ABCD Children's Report of Parental Behavioral Inventory.  crpbi01 |
| crpbi_parent3_y | First caregiver (caregiver participating in study/completing protocol). Is able to make me feel better when I am upset.  1 = Not like him/her; 2 = Somewhat like him/her; 3 = A lot like him/her | 1 = Not like him/her;  0 = otherwise | ABCD Children's Report of Parental Behavioral Inventory.  crpbi01 |
| crpbi_parent4_y | First caregiver (caregiver participating in study/completing protocol). Believes in showing his/her love for me.  1 = Not like him/her; 2 = Somewhat like him/her; 3 = A lot like him/her | 1 = Not like him/her;  0 = otherwise | ABCD Children's Report of Parental Behavioral Inventory.  crpbi01 |
| crpbi_parent5_y | First caregiver (caregiver participating in study/completing protocol). Is easy to talk to.  1 = Not like him/her; 2 = Somewhat like him/her; 3 = A lot like him/her | 1 = Not like him/her;  0 = otherwise | ABCD Children's Report of Parental Behavioral Inventory.  crpbi01 |
| crpbi_caregiver12_y | Second caregiver. Makes me feel better after talking over my worries with them.  1=Not like them; 2=Somewhat like them; 3=A lot like them | 1 = Not like him/her;  0 = otherwise | ABCD Children's Report of Parental Behavioral Inventory.  crpbi01 |
| crpbi_caregiver13_y | Second caregiver. Smiles at me very often.  1=Not like them; 2=Somewhat like them; 3=A lot like them | 1 = Not like him/her;  0 = otherwise | ABCD Children's Report of Parental Behavioral Inventory.  crpbi01 |
| crpbi_caregiver14_y | Second caregiver. Is able to make me feel better when I am upset.  1=Not like them; 2=Somewhat like them; 3=A lot like them | 1 = Not like him/her;  0 = otherwise | ABCD Children's Report of Parental Behavioral Inventory.  crpbi01 |
| crpbi_caregiver15_y | Second caregiver. Believes in showing their love for me.  1=Not like them; 2=Somewhat like them; 3=A lot like them | 1 = Not like him/her;  0 = otherwise | ABCD Children's Report of Parental Behavioral Inventory.  crpbi01 |
| crpbi_caregiver16_y | Second caregiver. Is easy to talk to.  1=Not like them; 2=Somewhat like them; 3=A lot like them | 1 = Not like him/her;  0 = otherwise | ABCD Children's Report of Parental Behavioral Inventory.  crpbi01 |
| ***Physical neglect***  *[To contribute a point to the score of Interpersonal Early Adversity (Family), at least one response should be coded as ‘1’]* | | | |
| parent_monitor_q1_y | How often do your parents/guardians know where you are?  1 = Never; 2 = Almost Never; 3 = Sometimes; 4 = Often; 5 = Always or Almost Always | 1 = Never/Almost Never;  0 = otherwise | ABCD Parental Monitoring Survey.  pmq01 |
| parent_monitor_q2_y | How often do your parents know who you are with when you are not at school and away from home?  1 = Never; 2 = Almost Never; 3 = Sometimes; 4 = Often; 5 = Always or Almost Always | 1 = Never/Almost Never;  0 = otherwise | ABCD Parental Monitoring Survey.  pmq01 |
| parent_monitor_q3_y | If you are at home when your parents or guardians are not, how often do you know how to get in touch with them?  1 = Never; 2 = Almost Never; 3 = Sometimes; 4 = Often; 5 = Always or Almost Always | 1 = Never/Almost Never;  0 = otherwise | ABCD Parental Monitoring Survey.  pmq01 |
| **Non-Familial Interpersonal Adversity (Range: 0-10)** | | | |
| ***Emotional abuse***  *[A ‘1’ on this item contributed a point to the score of Interpersonal Early Adversity (Non-Family)]* | | | |
| ksads_ptsd_raw_764_p | A non-family member threatened to kill your child.  0 = No; 1 = Yes | - | ABCD Parent Diagnostic Interview for DSM-5 (KSADS) Traumatic Events.  abcd_ptsd01 |
| ***Physical abuse***  *[A ‘1’ on this item contributed a point to the score of Interpersonal Early Adversity (Non-Family)]* | | | |
| ksads_ptsd_raw_761_p | Shot, stabbed, or beaten brutally by a non-family member.  0 = No; 1 = Yes | - | ABCD Parent Diagnostic Interview for DSM-5 (KSADS) Traumatic Events.  abcd_ptsd01 |
| ***Sexual abuse***  *[A ‘1’ on either item contributed a point to the score of Interpersonal Early Adversity (Non-Family)]* | | | |
| ksads_ptsd_raw_768_p | An adult outside your family touched your child in their privates, had your child touch their privates or did other sexual things to your child.  0 = No; 1 = Yes | - | ABCD Parent Diagnostic Interview for DSM-5 (KSADS) Traumatic Events.  abcd_ptsd01 |
| ksads_ptsd_raw_769_p | A peer forced your child to do something sexually.  0 = No; 1 = Yes | - | ABCD Parent Diagnostic Interview for DSM-5 (KSADS) Traumatic Events.  abcd_ptsd01 |
| ***Bullying***  *[A ‘1’ on this item contributed a point to the score of Interpersonal Early Adversity (Non-Family)]* | | | |
| kbi_p_c_bully | Does your child have any problems with bullying at school or in your neighborhood?  0 = No; 1 = Yes | - | ABCD Parent Diagnostic Interview for DSM-5 Background Items Full (KSADS-5)  dibf01 |
| ***School Safety***  *[A ‘1’ on this item contributed a point to the score of Interpersonal Early Adversity (Non-Family)]* | | | |
| school_6_y | I feel safe at my school.  1 = NO!; 2 = no; 3 = yes; 4 = YES! | 1 = NO!/no; 0 = otherwise | ABCD School Risk and Protective Factors Survey.  srpf01 |
| ***Other Community Violence***  *[A ‘1’ on any item contributed a point to the score of Interpersonal Early Adversity (Non-Family)]* | | | |
| ksads_ptsd_raw_758_p | Witnessed or present during an act of terrorism (e.g., Boston marathon bombing).  0 = No; 1 = Yes | - | ABCD Parent Diagnostic Interview for DSM-5 (KSADS) Traumatic Events.  abcd_ptsd01 |
| ksads_ptsd_raw_760_p | Witnessed someone shot or stabbed in the community.  0 = No; 1 = Yes | - | ABCD Parent Diagnostic Interview for DSM-5 (KSADS) Traumatic Events.  abcd_ptsd01 |
| neighborhood3r_p | My neighborhood is safe from crime.  1 = Strongly Disagree; 2 = Disagree; 3 = Neutral (neither agree nor disagree); 4 = Agree; 5 = Strongly Agree | 1 = Strongly Disagree/ Disagree;  0 = otherwise | ABCD Youth Neighborhood Safety/Crime Survey Modified from PhenX.  abcd_nsc01 |
| ***War***  *[A ‘1’ on this item contributed a point to the score of Non-Interpersonal Early Adversity]* | | | |
| ksads_ptsd_raw_759_p | Witnessed death or mass destruction in a war zone.  0 = No; 1 = Yes | - | ABCD Parent Diagnostic Interview for DSM-5 (KSADS) Traumatic Events.  abcd_ptsd01 |
| **Non-Interpersonal Adversity (Range: 0-5)** | | | |
| ***Natural Disaster***  *[A ‘1’ on this item contributed a point to the score of Non-Interpersonal Early Adversity]* | | | |
| ksads_ptsd_raw_757_p | Witnessed or caught in a natural disaster that caused significant property damage or personal injury.  0 = No; 1 = Yes | - | ABCD Parent Diagnostic Interview for DSM-5 (KSADS) Traumatic Events.  abcd_ptsd01 |
| ***Accident***  *[A ‘1’ on any item contributed a point to the score of Non-Interpersonal Early Adversity]* | | | |
| ksads_ptsd_raw_754_p | A car accident in which your child or another person in the car was hurt bad enough to require medical attention.  0 = No; 1 = Yes | - | ABCD Parent Diagnostic Interview for DSM-5 (KSADS) Traumatic Events.  abcd_ptsd01 |
| ksads_ptsd_raw_755_p | Another significant accident for which your child needed specialized and intensive medical treatment.  0 = No; 1 = Yes | - | ABCD Parent Diagnostic Interview for DSM-5 (KSADS) Traumatic Events.  abcd_ptsd01 |
| ksads_ptsd_raw_756_p | Witnessed or caught in a fire that caused significant property damage or personal injury.  0 = No; 1 = Yes | - | ABCD Parent Diagnostic Interview for DSM-5 (KSADS) Traumatic Events.  abcd_ptsd01 |
| ***Sudden Loss***  *[A ‘1’ on this item contributed a point to the score of Non-Interpersonal Early Adversity]* | | | |
| ksads_ptsd_raw_770_p | Learned about the sudden unexpected death of a loved one.  0 = No; 1 = Yes | - | ABCD Parent Diagnostic Interview for DSM-5 (KSADS) Traumatic Events.  abcd_ptsd01 |

**Table S1. Coding of early life adversity.**

### Supplementary Figures

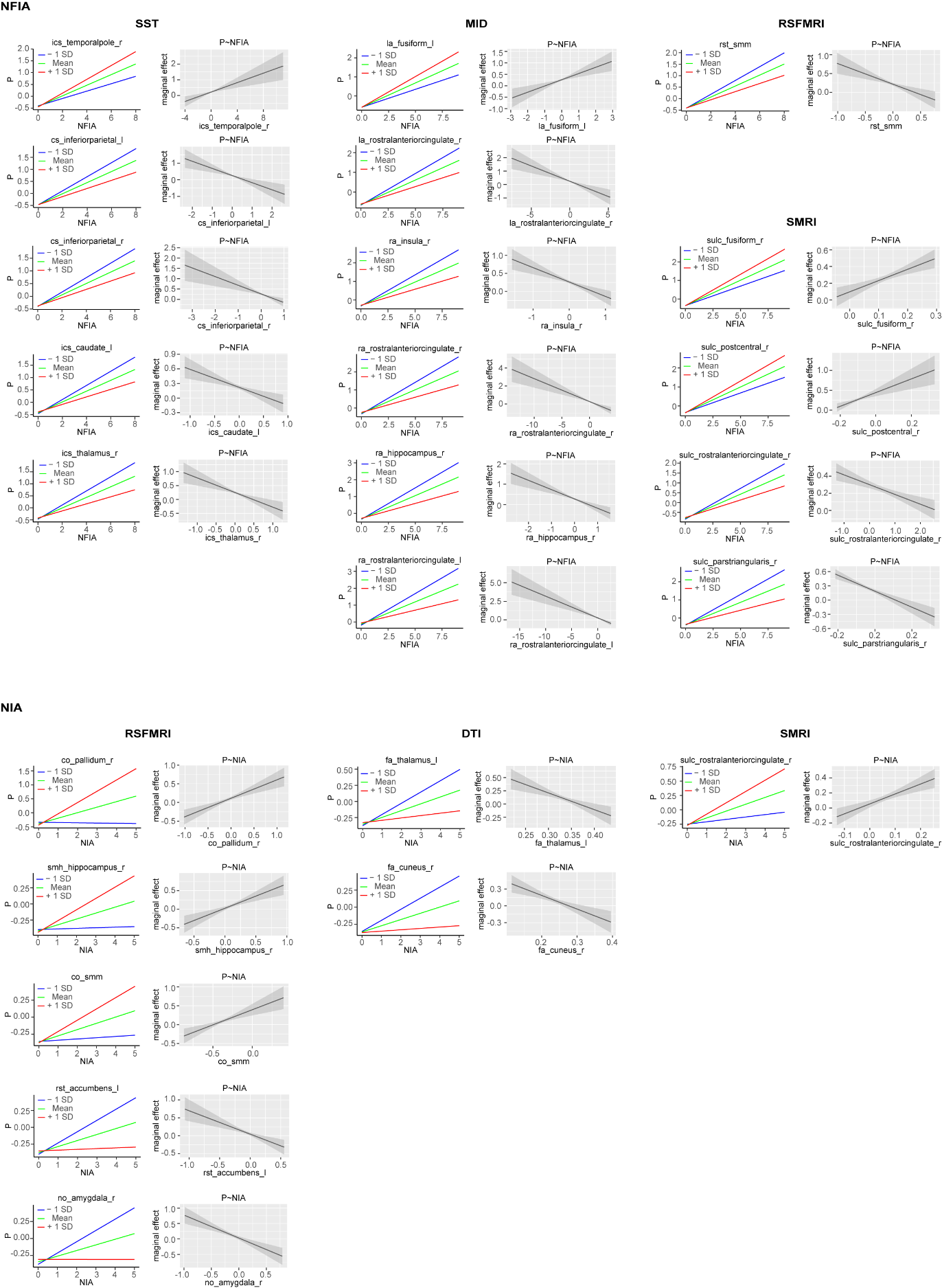

**Fig. S1: Simple slope and marginal effects analyses for brain features that significantly moderated the effect of ELA on P.**

Simple slope plots depict the ELA–psychopathology association at low, medium, and high levels of each brain feature. Marginal effects plots illustrate how the conditional effect of ELA on psychopathology varies across the continuous range of the brain feature, with shaded areas representing 95% CI.

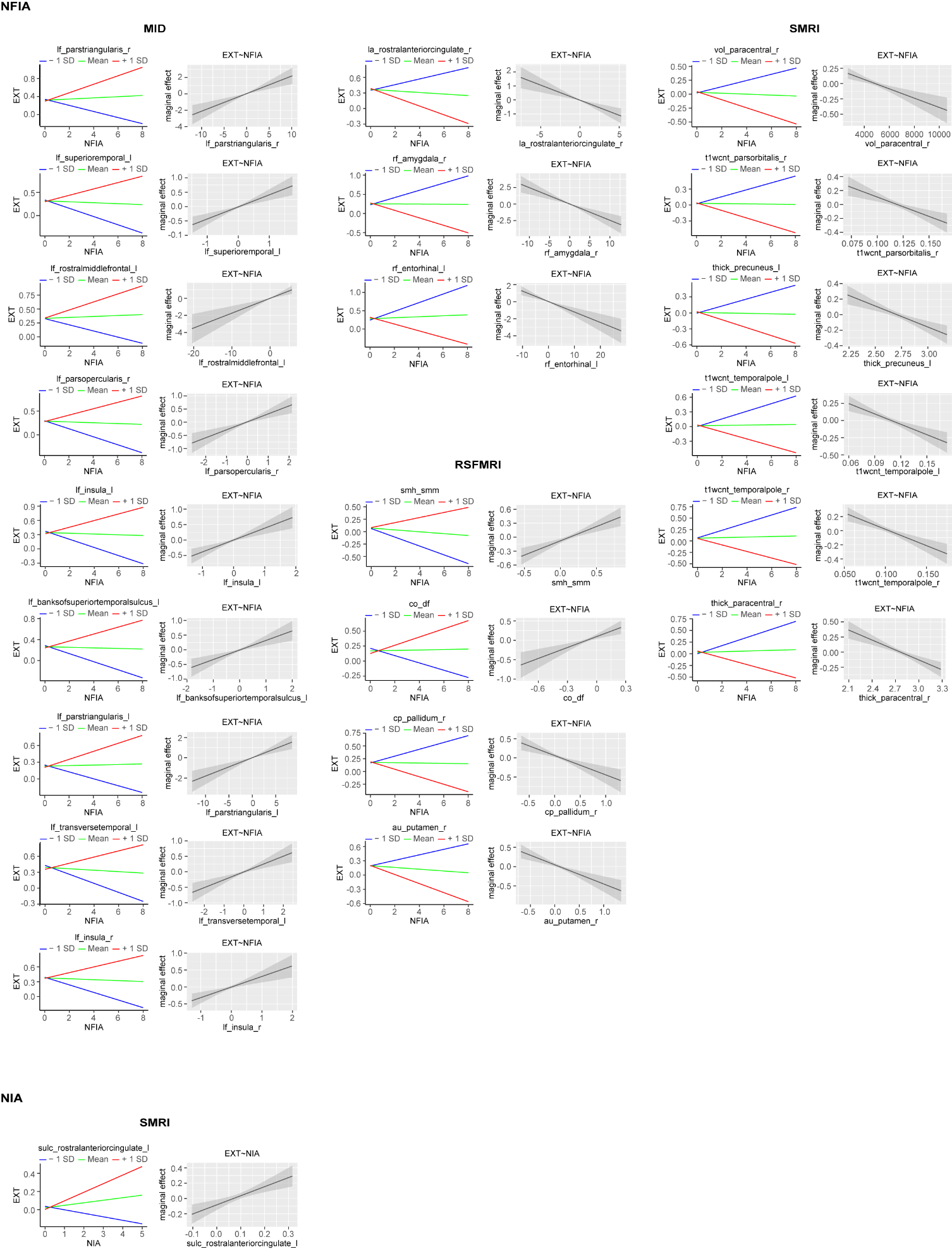

**Fig. S2: Simple slope and marginal effects analyses for brain features that significantly moderated the effect of ELA on EXT.**

Simple slope plots depict the ELA–psychopathology association at low, medium, and high levels of each brain feature. Marginal effects plots illustrate how the conditional effect of ELA on psychopathology varies across the continuous range of the brain feature, with shaded areas representing 95% CI.

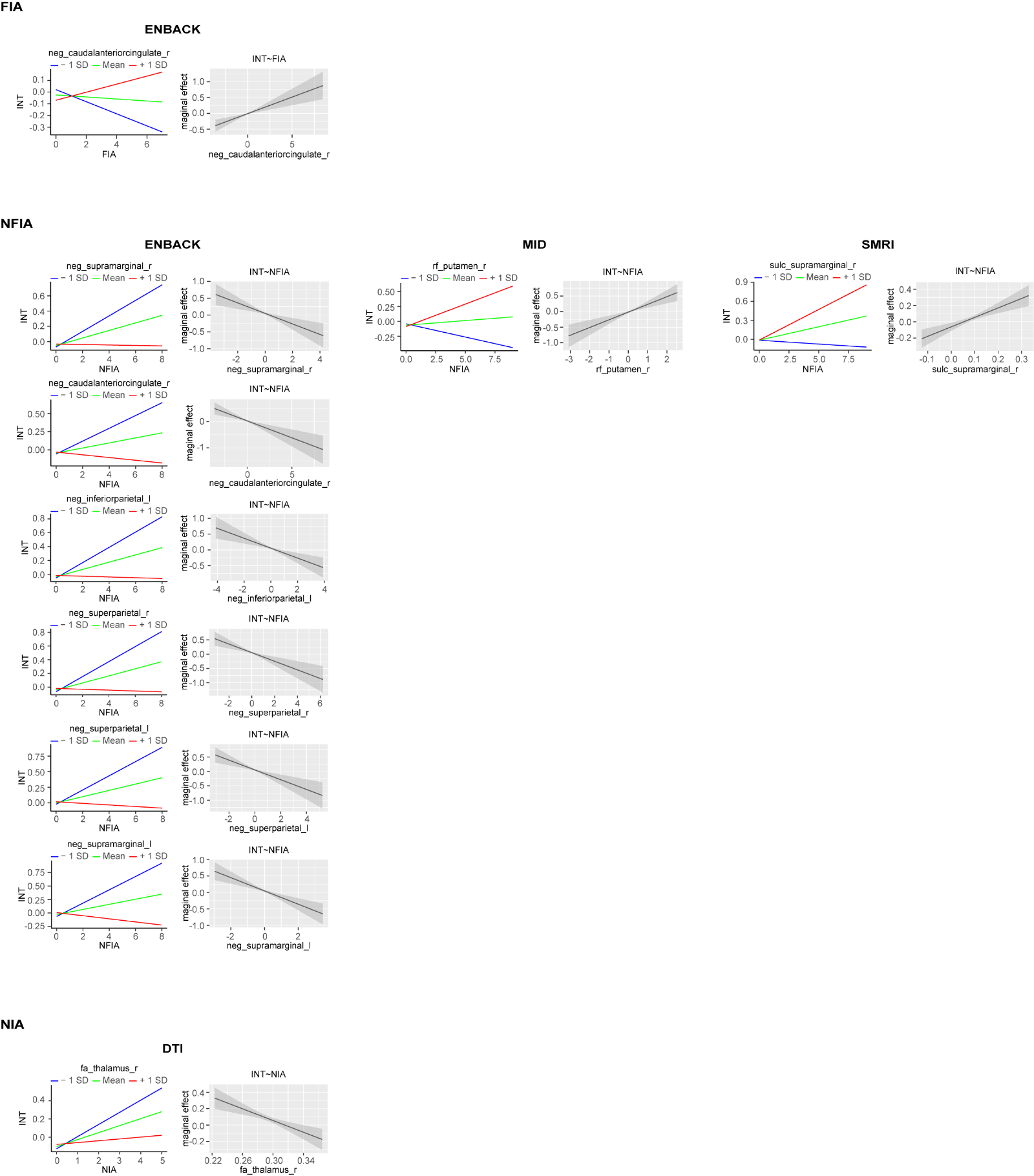

**Fig. S3: Simple slope and marginal effects analyses for brain features that significantly moderated the effect of ELA on INT.**

Simple slope plots depict the ELA–psychopathology association at low, medium, and high levels of each brain feature. Marginal effects plots illustrate how the conditional effect of ELA on psychopathology varies across the continuous range of the brain feature, with shaded areas representing 95% CI.
